# Smad6-mediated inhibition of BMP/TGF-β signaling disrupts midbrain growth in chick embryos

**DOI:** 10.64898/2026.03.30.714515

**Authors:** Dionysia Moschou, Anette Richter, Andrea Wizenmann

**Affiliations:** Department of Anatomie, Institute for Clinical Anatomy and Cell Analysis, Universität Tübingen, Germany; Department of Neurology and Neurosurgery, Centre for Research in Neuroscience, Research Institute of the McGill University Health Center, Montréal, Quebec H3G 1A4, Canada

## Abstract

Bone morphogenetic proteins (BMPs) play an important role in dorsal spinal cord patterning. Their presence in the roof plate of the midbrain indicates a role in its development. We examined whether the BMP signaling contributes to dorsal midbrain size expansion in chick embryos by missexpressing pathway activators and inhibitors. Overactivation of BMP4 did not affect midbrain development, whereas GDF7 reduced midbrain growth. In contrast, expression of a truncated dominant-negative BMP receptor type 1b or the extracellular inhibitor Chordin had no detectable effect. Ectopic expression of SMAD6, the intracellular inhibitor of the BMP/ TGF-β pathway, significantly reduced midbrain size, which correlated with decreased proliferation rates of SMAD6-overexpressing cells. In some cases, SMAD6 also disrupted MTN axon trajectory. These results indicate an important role for SMAD-dependent signaling pathways in early dorsal midbrain growth.

## Introduction

During development, morphogens pattern the dorsal-ventral axis of the neural tube. Sonic hedgehog (Shh), secreted by the floor plate and notochord, is important for ventral development in all brain regions together with FGF signaling in the forebrain *(Dessaud et al., 2007; Sagner & Briscoe, 2019; Watanabe & Nakamura, 2000)*.

Ventral specification promotes the generation of motor neurons and ventral interneurons in the spinal cord. In contrast, BMP (bone morphogenetic protein) and WNT proteins, which are highly expressed by the roof plate, play central roles in dorsal spinal cord patterning *(Chizikov & Millen, 2004; Wilson & Maden, 2005; Timmer et al., 2002)*, including the generation and specification of dorsal interneuron populations *(Liem et al., 1995; Lee et al., 1998; Butler & Dodd, 2003)*. These morphogens are also expressed in the anterior neural tube, which forms distinct brain regions *(Bobak et al., 2009; Bothe et al., 2011; GEISHA database)*, suggesting that BMP signaling may also be important for dorsal midbrain development.

### BMP pathway

BMPs were first discovered by their ability to direct the formation of bone and cartilage *(Wozney et al., 1988)*, but they are also required for the development of the dorsal spinal cord *(Liem et al., 1995; Lee et al., 1998)*, limb *(Bandyopadhyay et al., 2006)*, forebrain *(Hébert et al., 2002)*, eyes and kidneys *(Furuta & Hogan, 1998; Dudley et al., 1995)*. BMPs have a regulatory role in processes such as cell division, apoptosis, proliferation, cell migration, cell survival, and differentiation *(Chen et al., 2004)*. They belong to the TGF-β superfamily which includes more than 30 components such as TGF-βs, activins, osteogenic proteins (OPs), cartilage-derived morphogenetic proteins (CDMPs), or growth and differentiation factors (GDFs), all important for fate specification of neural tissue during development *(Ducy & Karsenty, 2000; Xiao et al., 2007)*. The importance of roof plate-derived BMP signals was shown by genetic ablation of the roof plate in spinal cord development *in vitro* and *in vivo (Millonig et al., 2000; Lee et al., 2000)*, ectopic ligand expression in the chick neural tube *(Lee et al., 1998; Liem et al., 1995; 1997)*, studies in mice and zebrafish lacking BMP receptors (BmpR) *(Millonig et al., 2000; Wine-lee et al., 2004; Lee et al., 200*0; *Nguyen et al., 2000)*, disrupted signaling *(Chesnutt et al., 2004)*, and siRNA-mediated knock-downs of BmpR signaling in chick embryos (Bobak et al 2009).

The extracellular signaling pathway includes BMP ligands, BMP antagonists, and proteinases that inactivate the BMP antagonists. Signal transduction is mediated by two serine-threonine (Ser/Thr) kinase receptors which heterodimerize upon ligand binding. BMP ligands bind first to the type 2 receptor (BmpR2) which phosphorylates the type 1 receptor (BmpR1), leading to phosphorylation of activated Smads (R-Smads). Two BmpR1 forms are expressed in the developing spinal cord, BmpR1a (ALK3) and BmpR1b (ALK6) *(Wine-Lee et al., 2004)*. SMAD1, SMAD5, and SMAD8 form a complex with SMAD4, translocate into the nucleus and regulate gene expression *(Kitisin et al, 2007)*.

Negative regulation occurs extracellularly through BMP antagonists that prevent ligand-receptor interaction, including the Cerberus family (including Grem1), Noggin, Chordin, and Follistatin *(Piccolo et al., 1996; Vonica & Brivanlou, 2006; Brazil et al., 2015; Dréau & Martí, 2013)*. The BMP and Activin Membrane Bound Inhibitor (BAMBI) acts as a pseudoreceptor at the cell membrane, reducing productive ligand-receptor signaling and activation of the BMP cascade *(Itoh & ten Dijke, 2007; Nohe et al., 2004)*. Receptor-mediated phosphorylation of Smad1/5/8 can be inhibited by FK-binding protein 12, inhibitory Smads (Smad6 and Smad7, i.e. I-Smads), ubiquitin ligase Smurf1 and cytosolic phosphatases, promoting degradation of BMP receptors or R-SMADs *(Hegarty et al., 2013)*. Smad6 further interferes with heteromeric complex formation by competing with Smad4 *(Hata et al. 1998; Nohe et al., 2004)*.

Importantly, Smad6/7 can also interact with components of the TGF-β pathway, leading to inhibition of Smad-dependent signaling more broadly. Together, numerous proteins regulate BMP signaling at multiple cellular levels.

### Midbrain development

Different neural structures develop from the midbrain, the second vesicle of the neural tube *(Watanabe & Yaginuma, 2015)*. In ventral midbrain, dopaminergic neurons of the substantia nigra and ventral tegmental area arise, together with oculomotor and trochlear motor neurons controlling eye movements *(Smidt & Burbach, 2007)*. In the early dorsal midbrain, the first neurons to differentiate form the mesencephalic trigeminal nucleus (MTN). Later, the superior (i.e. optic tectum) and inferior colliculi arise from the dorsal midbrain along with the pretectal nuclei *(Fedtsova & Turner, 2001; Gray & Sanes, 1992)*. The dorsal midbrain in chick consists mainly of the optic tectum, which extends more prominently than the ventral midbrain. The size difference is evident from embryonic day 3, when the dorsal midbrain spreads more laterally than the ventral midbrain (Figure 1). Dorsal midbrain cells undergo multiple rounds of proliferation to form the tecta *(Watanabe & Yaginuma, 2015)*. At this time point, ventral and dorsal midbrain identities are specified *(Li et al., 2005)*.

**Figure 1.**
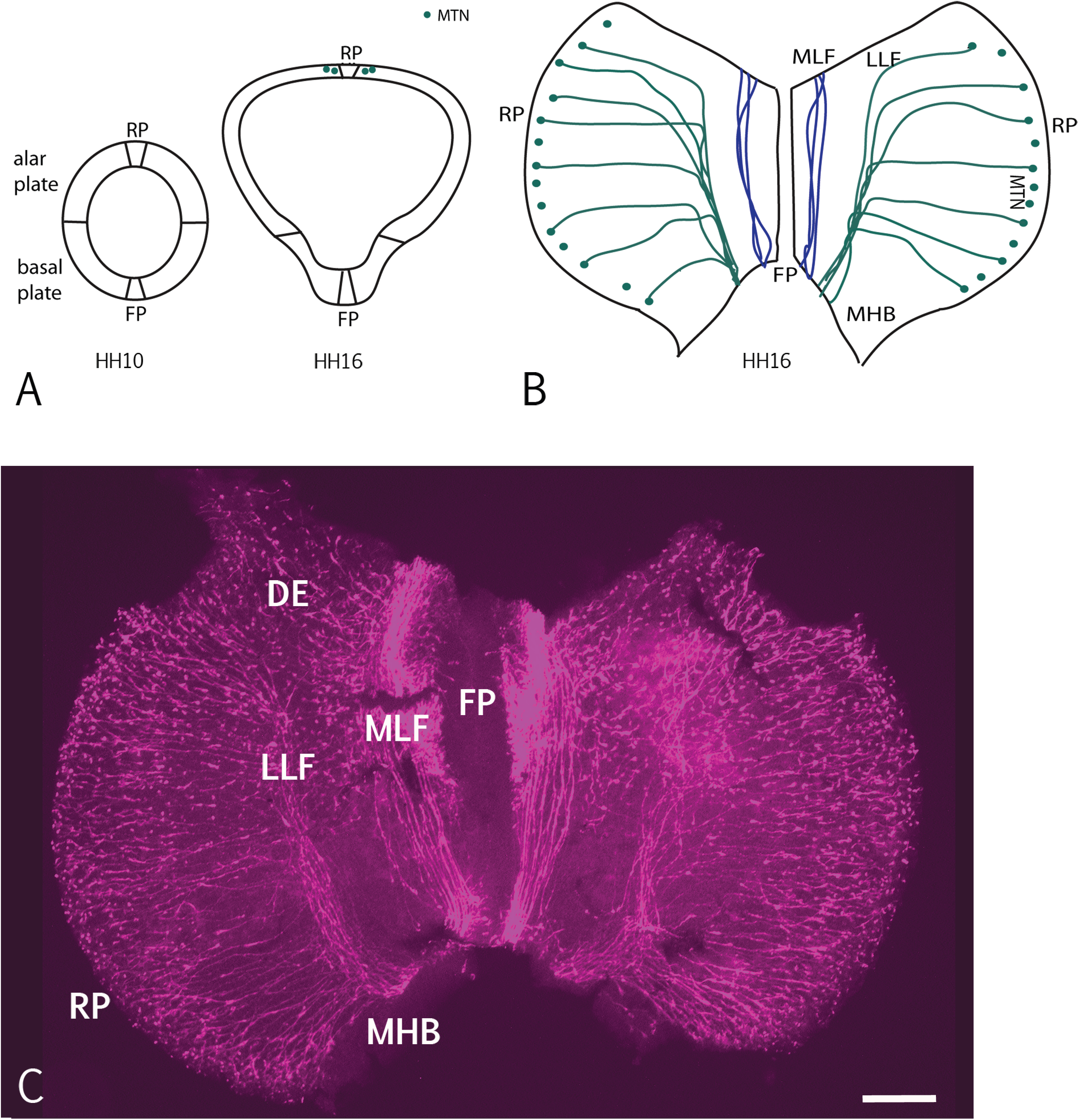
Schematics of the developing mesencephalon. (A) Schematics of transverse sections through the midbrain at HH10 and HH16. Alar plate, basal plate, roof plate, and floorplate are indicated. At HH10, the alar and basal plate show the same lateral extension. Two days later, at HH16, the alar plate extends more laterally than the basal plate. At HH16, neurons of the mesencephalic trigeminal nucleus (MTN; green dots, dorsal) are visible. (B) Schematic flat mount of an HH16 midbrain. MTN neurons on either side of the roof plate and the trajectory of their axons are indicated. The axons initially grow ventrally away from the roof plate. At a distance from the floor plate, they turn posteriorly towards the midbrain-hindbrain boundary (MHB) to form the lateral longitudinal fascicle (LLF). Axons from forebrain neurons form the more medially located medial longitudinal fascicle (MLF). (C) Flat-mounted HH16 midbrain stained for neurofilament using the 3A10 antibody (red) showing MTN axons and longitudinal fascicles. Schematics in (A) and (B) are not to scale. Scale bar (C): 100 µm. Abbr.: FP - floor plate, LLF – lateral longitudinal fascicle, MLF – Medial longitudinal fascile, MTN - mesencephalic trigeminal nucleus, RP - roof plate, DE – diencephalon.

BMPs are expressed in dorsal domains of the developing midbrain. Several BMP ligands (BMP4, BMP7, BMP8) and BMP receptors are expressed in the roof and alar plate of the chick midbrain vesicle *(GEISHA database; Bobak et al, 2009; Bothe et al., 2011)*. Of the two BmpR1 receptors, BmpR1b appears to be preferentially expressed in the midbrain vesicle *(Bobak et al, 2009)*. Interestingly, activation or blockade of BmpR1 signaling in chick midbrain did not change the expression of dorsal markers like PAX3, PAX7, MEIS2, or EFNB1 in dorsal midbrain *(Bobak et al. 2009)*, and deactivation of BMP signaling has no effect on the development of MTN neurons *(Lipovsek et al., 2017)*. Together, these findings suggest that BMP signaling plays a less prominent role in early fate specification in the dorsal midbrain *(Hébert et al., 2002; Watanabe & Yaginuma, 2015)* than in dorsal spinal cord patterning *(Liem et al., 1995; Lee et al., 1998; Butler & Dodd, 2003)*. This points to additional roles for BMP signaling during dorsal midbrain development.

### Influence of BMP signaling on cell proliferation

In addition to fate specification, BMP signaling has been implicated in the regulation of cell proliferation in a context-dependent manner. The influence of BMP signaling on cell proliferation depends on the cell type, the developmental stage and the BMP concentration in the tissue *(Brazil et al., 2015)*. Accordingly, BMP signal can either inhibit or promote cell cycle progression. In stem cells, it is known to regulate cell proliferation, differentiation, and apoptosis during embryogenesis *(Martinez et al., 2015)*. In many epithelial and stem cell systems, BMP signaling inhibits proliferation by inducing cell-cycle inhibitors (e.g. p21, p27) and repressing c-MYC, thereby promoting cell-cycle exit and differentiation *(Miyazono et al., 2010; Bhatia et al., 1999)*.

BMP signaling can also stimulate proliferation. An example of BMP enhancing proliferation is found in intestinal epithelium where it activates PI3K/AKT and thus leads to cell cycle progression *(Haramis et al., 2004; Auclair et al., 2007)*. In mesenchymal cells, osteoblasts, chondrocytes, and certain cancer cells, BMPs can stimulate proliferation, often via non-SMAD signaling and induction of Cyclin D1 and c-MYC, which facilitates G1/S progression *(Miyazono et al., 2010)*.

SMAD signaling pathways also control the expression of cell-cycle regulators such as p21, p27 (CDK inhibitors) and c-Myc, which in turn determine whether cells enter mitosis. A study by Fujita et al. *(2008)* showed that the loss of SMAD3 leads to an increased number of phospho-histone H3-positive mitotic cells in bone marrow stromal cells, indicating altered progression through mitosis. Furthermore, BMP/SMAD signaling pathway regulates mitotic checkpoint proteins, including BUB3 and MAD2, in breast cancer cells *(Yan et al., 2012)*. TGF-β/SMAD signaling can reduce mitosis by up-regulating CDK inhibitors such as p15INK4b and p21 *(Seoane et al., 2001)*. Thus, by regulating mitotic progression and cell-cycle control, BMP signaling may influence tissue growth, including the size of the dorsal midbrain.

Our study focuses on the effects of BMP signaling on dorsal midbrain growth in the chick embryo. Misexpression studies were conducted to assess the effects of BMP pathway overactivation or inhibition on midbrain development during early embryogenesis. Overexpressing the intracellular BMP/TGF-β inhibitor Smad6 in the developing dorsal midbrain, we observed growth deficits characterized by reduced midbrain size. Disrupting BMP signaling extracellularly by ectopic expression of a truncated dominant negative form of BMP receptor 1b did not replicate these growth effects. Notably, both forms of inhibition influenced MTN axon growth pattern.

## Results

### GFP expression did not influence midbrain growth

The chick midbrain is divided into the basal plate, which develops into the tegmentum, and the alar plate, which gives rise to the optic tecta (Fig. 1, schematic). The alar plate forming the tecta exhibits substantially higher proliferative activity than the basal plate. The distinction between dorsal (alar) and ventral (basal) midbrain territories is established early during development, as demonstrated by differential gene expression analyses in the chick midbrain by HH14–15 *(Chittka et al., 2009; Li et al., 2005)*.

We first tested whether the electroporation procedure itself influences the growth pattern or development of the transfected midbrain hemisphere. The dorsal midbrain was transfected with the vector pCAX expressing GFP (pCAX-GFP). Midbrains transfected with GFP and non-transfected (wildtype) were flat-mounted, imaged and analysed for the lateral extension. The relative sizes of left and right midbrain halves were calculated as a left to right area ratio. No difference in the size of midbrain hemispheres was observed between GFP-electroporated and wildtype samples (wildtype n = 5, area ratio = 0.99; GFP n = 3, area ratio = 1.00; p = 0.95; Fig. 2A,E). Therefore, we can conclude that electroporation with GFP alone had no measurable effect on midbrain growth.

**Figure 2.**
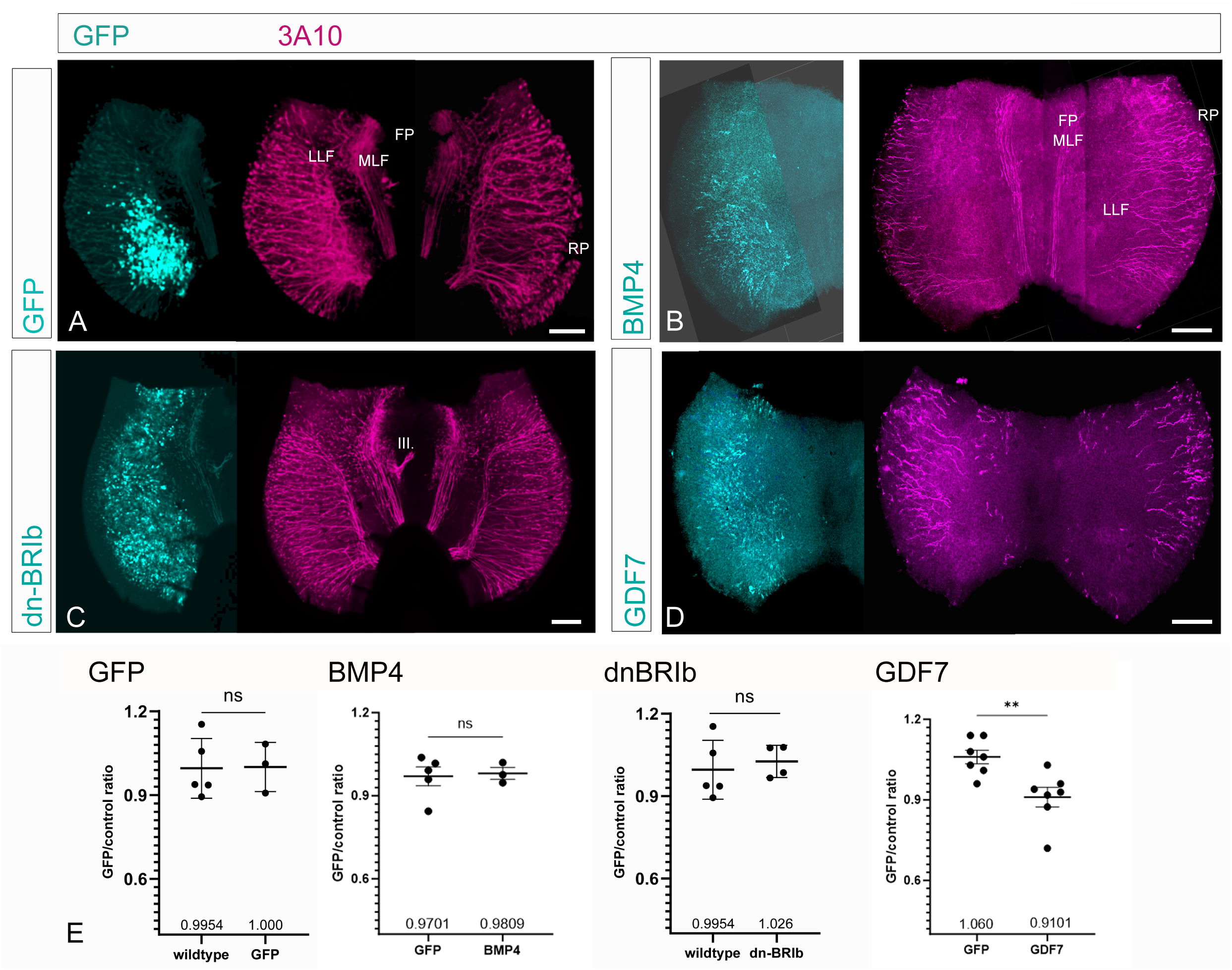
Differential effects of BMP pathway components on dorsal midbrain growth. Flat-mounted midbrains electroporated with GFP (A), BMP4 (B), dnBMPR1b (C), or GDF7 (D). The transfected cells express GFP (green). Axons and neurons are labeled in red using the 3A10 antibody and are shown on the right side of each panel (A-D). The transfected half of the midbrain is shown on the left side of each flat mount. The trajectories of the axons forming the LLF and MLF were not disrupted at transfected sites. (E) Quantification of midbrain size, expressed as the ratio of transfected to control sides. No significant difference in size was observed following GFP (p = 0.95), BMP4 (p = 0.79) and dnBR1b (p = 0.63) expression. In contrast, ectopic overexpression of GDF7 significantly decreased the midbrain size (p = 0.006). Comparison of means by Student’s t-test (mean ± SD are shown, *p < 0.05). Scale bar: 100 µm. Abbr.: III. – nervus oculomotorius, FP - floor plate, LLF – lateral longitudinal fascicle, MLF - medial longitudinal fascicles, RP -roof plate.

### Activation of the BMP pathway did not alter midbrain growth

Preliminary experiments in the laboratory indicated that BMP2-coated beads implanted into dorsal midbrain appeared to disturb MTN axon growth and potentially influence the size of the alar plate. To follow up on these observations, we interfered with intracellular and extracellular BMP signaling in the alar plate. We first ectopically expressed the BMP ligand BMP4 (Fig. 2B) and a constitutively active form of the BMPR1b receptor (ca-BR1b; data not shown) in the alar plate of the midbrain.

Quantification of the alar plate size showed that activation of the BMP signaling pathway by ectopic expression of BMP4 did not alter midbrain size (GFP control n = 5, area ratio = 0.97; BMP4 n = 3, area ratio = 0.98; p = 0.79; Fig. 2E). We also tested ectopic expression of ca-BR1B in midbrain however only in a single embryo (n = 1, area ratio = 0.95), which did not allow statistical analysis.

Inhibition of extracellular BMP signaling did not alter midbrain growth Next, we examined whether inhibition of the BMP signaling influences midbrain growth. The BmpR1b receptor is strongly expressed in the midbrain vesicle *(Bobak et al., 2009)*. Overexpression of the dominant-negative BMPR1b (dnBMPR1b) is expected to outcompete endogenous BmpR1b for dimerization with BMPR2 and thereby prevent signal transduction. We overexpressed a truncated, dominant-negative form of BmpR1b *(Bobak et al., 2009)* in the dorsal midbrain. Expression of dnBmpR1b did not affect midbrain growth. Statistical analysis indicated no significant effect of dnBmpR1b-mediated BMP inhibition on midbrain size (wildtype n = 5, area ratio = 0.99; dnBmpR1b n = 4, area ratio = 1.02; p = 0.63; Fig. 2C,E).

We also inhibited extracellular BMP signaling by ectopic expression of the BMP antagonist Chordin (CHRD; data not shown). Chordin is known to ventralize the neural tube *(Patten & Placzek, 2002)* and antagonize BMP signaling in the telencephalon *(Anderson et al., 2002; Piccolo et al., 1996)*. We also tested the effect of Chordin overexpression in one dorsal midbrain. In this midbrain growth was not altered, although a slight downward trend was observed (area ratio = 0.94).

### Overexpression of GDF7 reduced dorsal midbrain growth

We also overexpressed growth differentiation factor 7 (GDF7), a member of the TGF-β superfamily that is specifically expressed in the roof plate. GDF7 is involved in the induction of sensory neurons in dorsal spinal cord and restricts neural crest cells to a sensory fate *(Lee et al., 1998)*. GDF expression has also been reported in the dorsal midbrain of mice *(Lo et al., 2005)*.

Overexpression of GDF7 in the dorsal midbrain resulted in a significant reduction in the size of the transfected midbrain hemisphere compared to the control side (GFP control n = 7, area ratio = 1.06; GDF7 n = 7, area ratio = 0.91; p = 0.006; Fig. 2D,E). This indicates that elevated GDF7 levels negatively affect dorsal midbrain growth.

### Inhibition of intracellular BMP signaling influenced midbrain growth

Next, we ectopically expressed SMAD6, an intracellular inhibitor of BMP-Smad1/5/8 signal transduction. Overexpression of SMAD6 in the dorsal midbrain resulted in a reduced size of the transfected midbrain hemisphere compared to the control side (Fig. 3A,A’). The mean area ratio between the transfected and control sides was 0.82 ± 0.13 (n = 6). Statistical analysis using a Student’s t-test showed that this reduction was significant (p = 0.04; Fig. 3C). These results indicate that SMAD6-dependent signaling contributes to dorsal midbrain growth.

**Figure 3.**
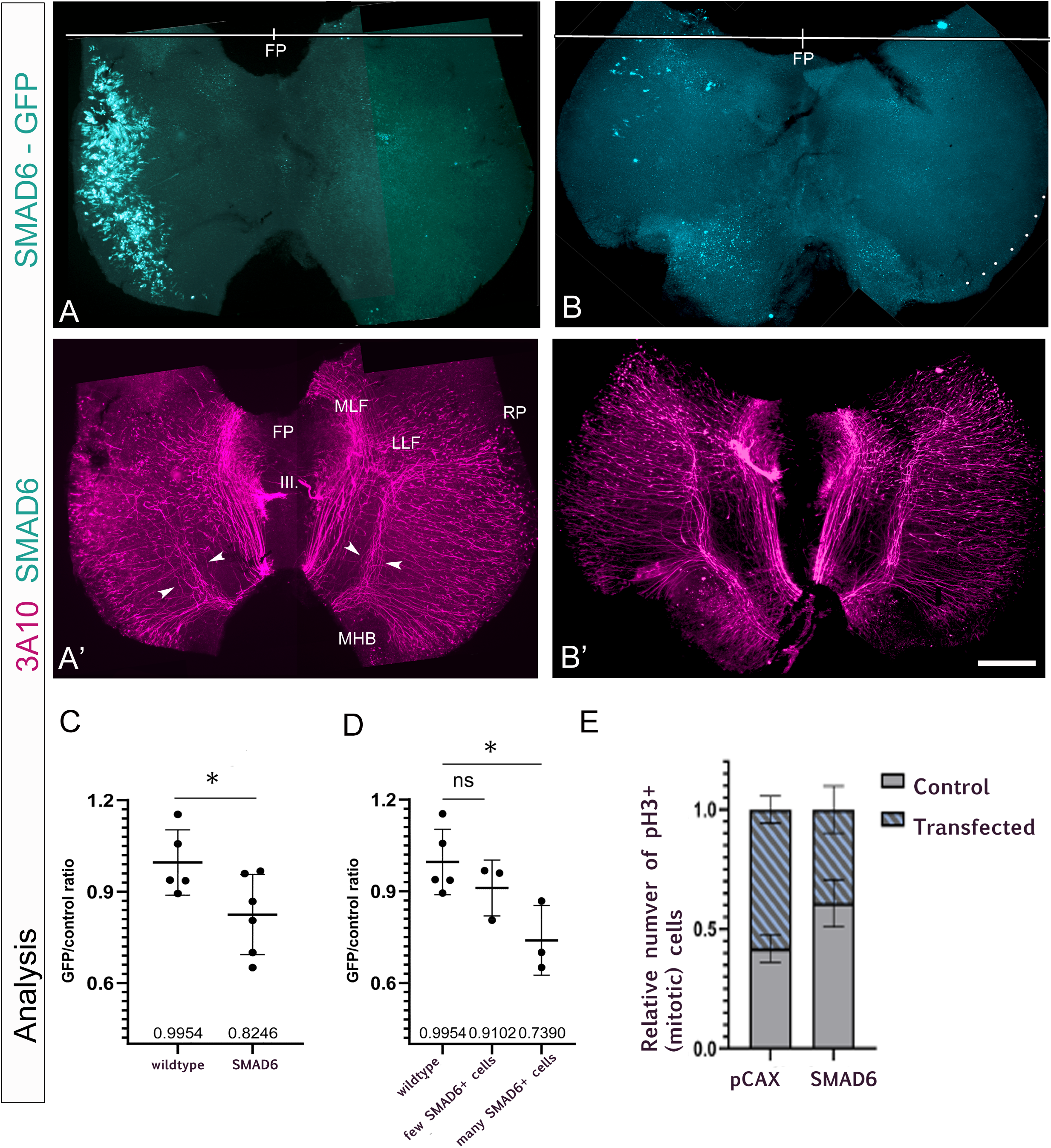
Smad6 reduces dorsal midbrain growth in a transfection-dependent manner. (A) SMAD6-GFP-labeled cells cover the dorsal left midbrain, which is smaller in size than the control side. The line at the top indicates the midline extension of both midbrain halves. (A’) MTN neurons and axons in the same flat mount are labeled using the 3A10 antibody. MTN axons grow toward the floor plate and turn posteriorly to form the LLF. Axons forming the LLF occupy a broader area on the transfected side compared to control (arrowheads). (B) Flat-mounted midbrain with sparse SMAD6-GFP-positive cells in the anterior left dorsal midbrain half. (B’) Neurons and axon trajectories are shown. The expression of SMAD6 has not affected the size of the transfected dorsal midbrain half and only minimally alters the LLF trajectory. (C-D) Quantification of midbrain size, expressed as the ratio of left to right halves. Strong SMAD6 expression reduces midbrain size. Comparison of means by Student’s t-test (mean ± SD are shown, *p < 0.05). (E) Quantification of PH3-positive cells in GFP- and SMAD6-transfected midbrain halves. GFP-transfected halves show no significant difference compared to control, whereas SMAD6-transfected halves show a significant reduction in pH3-positive cells. Comparison of means by two-way ANOVA (F(1,18) = 28.64, p<.001), mean ± SD are shown. Scale bar: 100 µm. Abbr: III. – nervus oculomotorius, FP - floor plate, LLF – lateral longitudinal fascicle, MLF - medial longitudinal fascicles, RP -roof plate.

Transfection efficiency varied between embryos (see Fig. 3A and 3B). We therefore examined whether the extent of Smad6 expression influenced midbrain growth.

When fewer cells were transfected (<50% of maximal transfection efficiency), SMAD6 overexpression did not result in a significant reduction in midbrain size (p = 0.26; n = 3; Fig. 3B,B’,C). In contrast, when a high proportion of cells (>50% of maximal efficiency) expressed SMAD6-GFP this reduction was significant (p = 0.01; n = 3; Fig. 3A,A’,C).

Although this study focused on the dorsal midbrain, we also assessed the effect of SMAD6 overexpression in the ventral midbrain in one example (n = 1). In this single case, SMAD6 expression in the ventromedial midbrain did not visibly affect midbrain size (data not shown), consistent with the predominantly dorsal expression of BMP signaling components.

### SMAD6 overexpression influenced cell proliferation

To examine the influence of SMAD6 on cell cycle progression, phospho-histone H3 (PH3) was used as a marker of mitosis, reflecting chromatin condensation and entry into M-phase. GFP or SMAD6 were ectopically expressed in one dorsal midbrain hemisphere, and the number of PH3-positive cells was compared between the electroporated and control sides.

Overexpression of GFP resulted in 59% of PH3-positive cells on the electroporated midbrain half, compared to 41% on the control side. In contrast, ectopic expression of SMAD6 produced the opposite effect, with 39% of PH3-positve cells on the Smad6-expressing side and 60% on the control side (Fig. 3D). Two-way ANOVA revealed a significant interaction between construct (GFP vs SMAD6) and electroporation status (control vs electroporated hemisphere) (F(1,18) = 28.64, p<.001). Post hoc Šídák comparisons confirmed that GFP electroporation increased the proportion of PH3-positive cells relative to the control side (p = 0.034, n = 5), whereas SMAD6 electroporation significantly reduced the proportion of PH3-positive cells (p = 0.002, n = 6). These results indicate that SMAD6 overexpression reduced the proportion of cells entering mitosis. This reduction correlates with the decreased size of dorsal midbrain hemispheres overexpressing SMAD6.

### Inhibition of BMP signaling disrupted MTN axonal pathfinding

Analysis of midbrain flat mounts transfected with SMAD6 revealed that, in several cases, the growth direction of mesencephalic trigeminal nucleus (MTN) axons was disrupted when transfection was broad (3 out of 11 embryos; Fig.3A’,4B,B’). In these embryos, MTN axons deviated from their typical ventral trajectory. These observations suggest that BMP signaling may contribute to MTN axon pathfinding.

Consistent with this finding, altered MTN axon trajectories were also observed in a subset of embryos following ectopic expression of dominant-negative BMPR1b, with 2 out of 7 embryos displaying MTN axons that failed to extend straight ventrally (Fig. 4A,A’). Midbrain halves transfected with GDF7 displayed a mostly normal lateral longitudinal fasciculus (LLF) course (1 out of 11 was disturbed). Although the penetrance of this phenotype was low and additional experiments will be required to quantify this effect, these results indicate that disruption of BMP signaling might affect MTN axon guidance.

**Figure 4.**
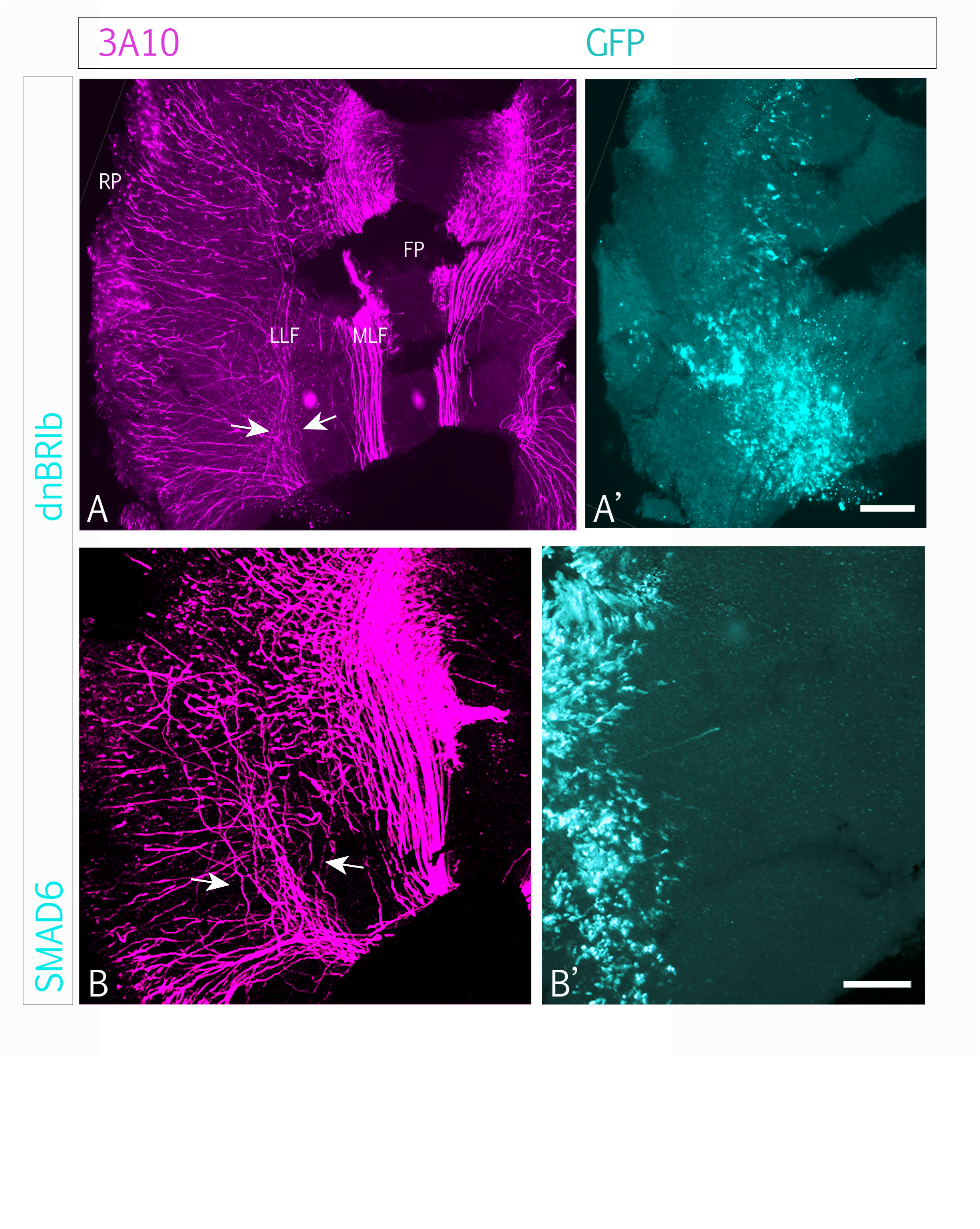
dnBR1b and Smad6 overexpression disrupts MTN axon trajectories in the dorsal midbrain. (A) Flat-mounted midbrain transfected with dnBMPR1b in the left half (A’). On the transfected side, formation of the LLF by MTN axons (arrows in A) is disrupted in regions with strong dnBMPR1b expression (compare to A’). (B) Higher magnification of SMAD6-transfected midbrain from Figure 3A,A’. The arrow indicates the broader trajectory of the LLF. Scale bar: 100 µm. Abbr.: FP - floor plate, LLF – lateral longitudinal fascicle, MLF - medial longitudinal fascicles, RP -roof plate.

## Discussion

The presence of multiple components of the BMP signaling pathway in the alar plate of the dorsal midbrain suggests a role for this pathway in dorsal midbrain development, including formation of the future optic tectum and mesencephalic trigeminal nucleus (MTN). However, previous studies have shown that experimental overactivation or suppression of BMP signaling does not alter the expression of alar plate-specific markers *(Bobak et al., 2009)* or the induction of MTN neurons *(Lipovsek et al. 2017)*.

In the present study, we show that disrupting intracellular BMP/TGF-β signaling with SMAD6 overexpression in the midbrain alar plate produces distinct outcomes depending on the level at which the pathway is inhibited. Overexpression of BMP4 did not influence midbrain growth. Interestingly, overexpression of GDF7 lead to a reduction in size of the transfected midbrain half. Interruption of extracellular BMP signaling using a truncated dominant-negative BMPR1b did not affect midbrain growth or MTN axon development. The inhibition of intracellular BMP signaling by SMAD6 overexpression at embryonic day 2 resulted in a significant reduction in the size of the transfected alar plate and in some cases in disturbances in MTN axon growth direction. The magnitude of the growth reduction correlated with the extent of transfection. Strong dorsal SMAD6 expression led to midbrain halves that were markedly smaller than the control side.

The lack of a detectable phenotype following dnBMPR1b expression, despite the pronounced effects of SMAD6, likely reflects redundancy within the BMP receptor system. In the developing dorsal midbrain, several BMP ligands, including BMP2, BMP4, and BMP7, are present and can signal through multiple type 1 receptors such as BMPR1A (ALK3), BMPR1B (ALK6) *(Wine-Lee et al., 2004)*, and in some contexts ACVR1 (ALK2) *(Miyazono et al., 2010)*. Blocking a single receptor subtype may therefore be insufficient to substantially reduce pathway activity, whereas SMAD6 inhibits intracellular BMP transduction downstream of all type 1 receptors by interfering with SMAD1 signaling and preventing formation of the active SMAD1-SMAD4 transcriptional complex *(Hata et al., 1998)*. Differences in construct efficiency, receptor stoichiometry, and temporal expression patterns may further contribute to the lack of effect observed with dnBMPR1B. The other possibility is that the TGF-β pathway is involved in the growth that is the proliferation rate of cells in the midbrain, Further investigations are necessary to assess the role of the TGF-β pathway.

Given these considerations, it is also possible that compensatory mechanisms preserve BMP signaling following dnBMPR1b expression. Assessing phospho-Smad1/5/8 (P-Smad1/5/8) levels in dnBMPR1b-electroporated tissue would help determine whether BMP signaling is effectively reduced. If P-Smad1/5/8 levels remain unchanged, this would suggest either compensation by other receptors or insufficient inhibition by the dominant-negative construct.

Our observations prompted us to consider how BMP signaling interacts with other pathways controlling proliferation and differentiation. In several contexts, removal of BMP signaling elevates WNT activity, leading to increased proliferation and reduced differentiation, as reported in the spinal cord *(Ille et al., 2007; Dréau & Martí, 2013)* and telencephalon *(Solloway & Robertson, 1999; Bachiller et al., 2000)*. Conversely, BMP overexpression promotes premature differentiation of neural progenitors in the spinal cord *(García-Campmany & Martí, 2007)*, an effect that is notably not observed in the dorsal midbrain *(Lipovsek et al., 2017)*. Together, these findings highlight the context-dependent nature of BMP signaling and suggest that its interaction with other pathways may regulate growth in the midbrain alar plate.

In this context, SMAD6-mediated inhibition likely affects not only BMP-Smad-dependent pathways. SMAD6 can interfere with signaling downstream of both BMP and TGF-β ligands *(Itoh & Dijke, 2007)*, indicating a complex intracellular network that may influence proliferation and growth of the dorsal midbrain. Candidate interacting pathways include WNTs/β-catenin signaling *(Quinlan et al., 2009)* and growth differentiation factor 7 (GDF7), which is expressed in the roof plate and has established roles in dorsal neural development *(Vonica & Brivanlou, 2006)*, as well as the TGF-β pathway. Indeed, TGF-β signaling has been implicated in the development of the dopaminergic and other neurons in the ventral midbrain *(Roussa et al., 2009; Hegarty et al., 2014; Brodski et al., 2019)*. This suggests that both BMP and TGF-β pathways may act complementary but in distinct ways across midbrain domains. Although TGF-β2 is the only TGF-β ligand reported in the early midbrain *(GEISHA database)*, the contribution of additional TGF-β superfamily components cannot be excluded.

The reduction in alar plate size following SMAD6 overexpression is consistent with decreased proliferation of neuronal progenitors. This interpretation is supported by our analysis of PH3-positive cells, which revealed reduced mitotic activity in SMAD6-transfected tissue. Alternative explanations, such as premature differentiation or increased apoptosis, cannot be ruled out and could be addressed in future studies using differentiation markers or apoptosis assays.

The dorsal midbrain undergoes extensive proliferation to form a swelling that subsequently separates into two hemispheres at the alar plate *(Watanabe & Yaginuma, 2015)*, as illustrated in Figure 1. Disruption of BMP-dependent signaling during this period may therefore impact both growth dynamics and axonal organization.

While our study focused on embryos at HH16-18, it remains possible that BMP signaling influences midbrain development at earlier or later stages. Temporal manipulation of BMP/TGF-β pathway activity may therefore reveal additional roles in midbrain growth or circuit formation.

In addition to growth effects, SMAD6 overexpression disrupted MTN axon trajectories. Consistent with this observation, preliminary experiments using BMP2-coated beads also interfered with MTN axon growth. Our data suggest that MTN axonal growth ventrally to form the LLF might be influenced by BMP signaling. We found that in some cases the formation of the LLF by MTN axons was partially disturbed upon strong overexpression of SMAD6 or dnBR1b and, in one case also by overexpression of GDF7.

These findings align with previous reports demonstrating that BMPs contribute to motor axon navigation *(Jardin et al., 2018)* and play broader roles in shaping axonal architecture *(Fassier et al., 2010; Tsang et al., 2009; Wang et al., 2007)*. In ventral midbrain, BMP signaling was shown to be essential for the differentiation of midbrain dopaminergic neurons *(Jovanovic et al., 2018; Hegarty et al., 2014)*. BMP2 has also been shown to promote neurite outgrowth in midbrain dopaminergic neurons, which share aspects of their developmental programs with MTN neurons *(Hegarty et al., 2014; O’Keefe et al., 2017)*. In spinal cord, WNT and BMP proteins produced by the roof plate, control proliferation, specification, migration and axon guidance of adjacent dorsal interneurons in the spinal cord *(Chizihkov & Millen, 2005)*. BMPs have been also shown that they can modulate neuronal responsiveness in axonal regeneration where they influence the directionality of axonal growth *(Zhong & Zou, 2014)*. A reason for the scattered growth pattern could be the influence of BMP signaling on the dorso-ventral determination of the cells. The induction of dorsally located proteins could induce disturbed axonal growth. Li et al, (2005) showed that the dorso-ventral axis of the midbrain is progressively specified. It can still be influenced up to HH17. Another reason for the disturbed axonal growth might be the reduced extension of the dorsal midbrain. This could have an effect on the distribution of chemokines guiding MTN axons ventrally and then posteriorly.

Additional experiments are required to further investigate the mechanisms and pathway interactions involved in midbrain growth and MTN axon guidance. This study highlights the importance of Smad-dependent, multifaceted signaling pathways during chick midbrain development.

Our findings indicate that extracellular manipulation of individual BMP receptors has limited impact on dorsal midbrain growth. The intracellular inhibition of Smad-dependent signaling however, significantly affected both tissue growth and in some cases the close formation of the LLF by MTN axons. SMAD6 overexpression profoundly altered midbrain development, reducing alar plate size, whereas blockade of a single BMP receptor subtype had little effect. These results highlight both the redundancy of receptor usage and the broader inhibitory scope of intracellular regulators like Smad6. Our findings suggest that BMP signaling in the dorsal midbrain is not directly important for midbrain growth whereas other components of the TGF-β superfamily might play an important role. Taken together our results suggest that SMAD6-dependent signaling emerges as a critical regulator of multiple developmental processes in the chick midbrain, extending beyond fate specification to include growth control and axonal organization.

## Materials and methods

### *In ovo* electroporation

Fertilized LB eggs were incubated at 37°C to the required Hamburger and Hamilton (HH), stage *(Hamburger and Hamilton, 1992)* for electroporation (HH9 to HH11) as previously described *(Huber et al., 2013; Itasaki et al., 1999, Momose et al., 1999)*. In short, 1-2 ml of albumin was removed to lower the embryo. A window was cut into the upper side of the egg with scissors. The midbrain neural tube was transfected with the different vectors at a concentration of 1 µg/µl. The used vector constructs with different BMP and/or GFP are listed in Table 1. Four to five 10 ms / 15 V pulses were applied by electrodes placed on either side of the midbrain. Then approximately 500 ml phosphate buffered saline (PBS) with antimycoticum and antibioticum was added before eggs were sealed. Eggs were incubated until embryos reached HH16 to HH18 and then fixed.

**Table 1.** Expression vectors and gene inserts.

| PLASMID | PLASMID LENGTH (kb) | INSERT | INSERT LENGTH (kb) |
| --- | --- | --- | --- |
| pCAX | 4.2 | GFP | 0.7 |
| pCDNA 3.1/myc-His | 5.5 | Bmp4 | 2 |
| pMES IRES GFP | 5.6 | dn-BR1b | 1.0 |
| pCAβ IRES GFP | 5.7 | SMAD6 | 1.3 |
| pMES IRES GFP | 5,7 | BR1b-ca | 1,6 |
| pCAβ IRES GFP | 5.7 | CHORDIN | 2.9 |

### Flat mounts of midbrain tissue

For flat mounts, the midbrain was separated from the rest of the embryo, and the surrounding mesenchyme was removed. A cut of the roof plate along its anterior posterior axis allowed the flattening of the tissue on a microscope slide with the basal side facing the cover slip. To embed the tissue, a mixture of PBS and Glycerol (50% each with 0.01% Triton X-100) was used.

### Cryosections of midbrain tissue

For cryosections embryos were incubated in 30% sucrose overnight. The heads were then transferred to a Cryomold and filled with the OCT compound (Tissue Tek) until the head was fully covered. To adapt to OCT, the head was left in the compound for at least a minute before it was frozen. To be able to perform coronal sections of the midbrain, the head was accordingly arranged and by placing the mold on dry ice (under a stereomicroscope) the head could be frozen with the required orientation. Consecutive sections of 20 μm in the direction from dorsal to ventral were cut with a cryotome (Leica) and collected with coated slides. The sections were dried overnight at 4°C and stored at -20°C.

### Immunofluorescence

Fixed, electroporated whole midbrains or cryosections were washed three times in blocking solution (PBS supplemented with 10% fetal calf serum (FCS) and 0.1% Triton X-100). For whole-mount staining, primary antibodies diluted in blocking solution were added and samples were incubated at 4°C for 2 to 3 days. Samples were then washed at least five times in PBS and incubated overnight at 4°C with secondary antibodies diluted in blocking solution.

Cryosections were incubated with primary antibodies overnight at 4°C, followed by three washes in PBS for 5 min and one wash in blocking solution for 30 min. Secondary antibodies were then applied for 2 h at room temperature. Sections were washed again and mounted in glycerol/PBS (1:9).

Primary antibodies used were 3A10 (neurofilament associated antibody; 3A10; Developmental Studies Hybridoma Bank, Iowa) to label neurons and axons. For cell cycle analysis, we used the anti-Phospho-Histone H3 (Ser10; PH3) to stain the late G2 beginning of M-phase. GFP was detected using a rabbit anti-GFP antibody (MoBiTec). Secondary goat anti-mouse Cy3 IgG and anti-rabbit Alexa A488 IgG were used at 1:500 dilution (Molecular Probes).

### Imaging, cell counting and statistical analysis

Z-stack images of immunostainings were taken with a ZEISS Observer.Z1 microscope with a ZEISS AxioCam MRc or a ZEISS LSM 510 invert Confocal Microscope with a ZEISS AxioCam MRc.

Images were analyzed using ImageJ (NIH). To quantify midbrain growth, we measured the two midbrain halves using ImageJ (in pixels), and we calculated the ratio of the electroporated side to the non-electroporated side. The size of area of the electroporated (GFP^+^) side was compared to the area of the control (GFP^−^) side by calculating their ratio (GFP/control). The GFP/control ratio for each introduced gene was compared to the left side/right side ratio of the non-electroporated (wildtype) samples.

To assess proliferation, PH3-positive cells were quantified in sections overexpressing GFP alone or SMAD6/GFP. For each condition, the proportion of PH3+ cells among GFP+ cells was calculated. At least two distinct regions per section were analyzed, and the mean and standard deviation were calculated.

Statistical analysis and graph design were performed using Prism 8 and 10 (Graphpad). The normality of the data distribution was assessed using the Shapiro-Wilk test. Two-tailed Student’s t-test was used to evaluate differences between two groups, with Welch’s correction applied when standard deviations of the groups differed. For experiments involving two independent variables, two-way ANOVA was performed followed by Šídák’s multiple comparisons test. A p-value of 0.05 was used as the threshold for statistical significance.

## Acknowledgments

We are grateful to Lothar Just and Anthony Graham for helpful discussions and comments on the manuscript.

## Author Contributions

Conceptualization, A.R., A.W.; Investigation, D.M., A.R., A.W.; Formal analysis, D.M., A.R.; Writing – original draft, D.M.; Writing – review and editing, D.M., A.W.; Supervision, A.W.; Funding acquisition, A.W.

## References

Anderson, R. M., Lawrence, A. R., Stottmann, R. W., Bachiller, D., & Klingensmith, J. (2002). Chordin and noggin promote organizing centers of forebrain development in the mouse. Development (Cambridge, England), 129(21), 4975–4987. 10.1242/dev.129.21.4975

Auclair, B. A., Benoit, Y. D., Rivard, N., Mishina, Y., & Perreault, N. (2007). Bone morphogenetic protein signaling is essential for terminal differentiation of the intestinal secretory cell lineage. Gastroenterology, 133(3), 887–896. 10.1053/j.gastro.2007.06.066

Bachiller, D., Klingensmith, J., Kemp, C., Belo, J. A., Anderson, R. M., May, S. R., McMahon, J. A., McMahon, A. P., Harland, R. M., Rossant, J., & De Robertis, E. M. (2000). The organizer factors Chordin and Noggin are required for mouse forebrain development. Nature, 403(6770), 658–661. 10.1038/35001072

Bandyopadhyay, A., Tsuji, K., Cox, K., Harfe, B. D., Rosen, V., & Tabin, C. J.. (2006). Genetic Analysis of the Roles of BMP2, BMP4, and BMP7 in Limb Patterning and Skeletogenesis. Plos Genetics, 2(12), e216. 10.1371/journal.pgen.0020216

Bhatia, M., Bonnet, D., Wu, D., Murdoch, B., Wrana, J., Gallacher, L., & Dick, J. E. (1999). Bone morphogenetic proteins regulate the developmental program of human hematopoietic stem cells. The Journal of experimental medicine, 189(7), 1139–1148. 10.1084/jem.189.7.1139

Bobak, N., Agoston, Z., & Schulte, D. (2009). Evidence against involvement of Bmp receptor 1b signaling in fate specification of the chick mesencephalic alar plate at HH16. Neuroscience letters, 461(3), 223–228. 10.1016/j.neulet.2009.06.003

Bothe, I., Tenin, G., Oseni, A., & Dietrich, S. (2011). Dynamic control of head mesoderm patterning. Development (Cambridge, England), 138(13), 2807–2821. 10.1242/dev.062737

Brazil, D. P., Church, R. H., Surae, S., Godson, C., & Martin, F. (2015). BMP signalling: agony and antagony in the family. Trends in cell biology, 25(5), 249–264. 10.1016/j.tcb.2014.12.004

Brodski, C., Blaess, S., Partanen, J., & Prakash, N. (2019). Crosstalk of Intercellular Signaling Pathways in the Generation of Midbrain Dopaminergic Neurons In Vivo and from Stem Cells. Journal of Developmental Biology, 7(1), 3. 10.3390/jdb7010003

Butler, S. J., & Dodd, J. (2003). A role for BMP heterodimers in roof plate-mediated repulsion of commissural axons. Neuron, 38(3), 389–401. 10.1016/s0896-6273(03)00254-x

Chen, D., Zhao, M., & Mundy, G. R. (2004). Bone Morphogenetic Proteins. Growth Factors, 22(4), 233–241. 10.1080/08977190412331279890

Chesnutt, C., Burrus, L. W., Brown, A. M., & Niswander, L. (2004). Coordinate regulation of neural tube patterning and proliferation by TGFbeta and WNT activity. Developmental biology, 274(2), 334–347. 10.1016/j.ydbio.2004.07.019

Chittka, A., Volff, J., & Wizenmann, A. (2009). Identification of genes differentially expressed in dorsal and ventral chick midbrain during early development. BMC developmental biology, 9, 29. 10.1186/1471-213X-9-29

Chizhikov, V. V., & Millen, K. J. (2004). Mechanisms of roof plate formation in the vertebrate CNS. Nature reviews. Neuroscience, 5(10), 808–812. 10.1038/nrn1520

Chizhikov, V. V., & Millen, K. J. (2005). Roof plate-dependent patterning of the vertebrate dorsal central nervous system. Developmental biology, 277(2), 287–295. 10.1016/j.ydbio.2004.10.011

Dessaud, E., Yang, L. L., Hill, K., Cox, B., Ulloa, F., Ribeiro, A., Mynett, A., Novitch, B. G., & Briscoe, J. (2007). Interpretation of the sonic hedgehog morphogen gradient by a temporal adaptation mechanism. Nature, 450(7170), 717–720. 10.1038/nature06347

Dréau, G. L. and Martí, E. (2013). The multiple activities of bmps during spinal cord development. Cellular and Molecular Life Sciences, 70(22), 4293–4305. 10.1007/s00018-013-1354-9

Ducy, P., & Karsenty, G. (2000). The family of bone morphogenetic proteins. Kidney international, 57(6), 2207–2214. 10.1046/j.1523-1755.2000.00081.x

Dudley, A. T., Lyons, K. M., & Robertson, E. J.. (1995). A requirement for bone morphogenetic protein-7 during development of the mammalian kidney and eye.. Genes & Development, 9(22), 2795–2807. 10.1101/gad.9.22.2795

Fassier, C., Hutt, J. A., Scholpp, S., Lumsden, A., Giros, B., Nothias, F., Schneider-Maunoury, S., Houart, C., & Hazan, J. (2010). Zebrafish atlastin controls motility and spinal motor axon architecture via inhibition of the BMP pathway. Nature neuroscience, 13(11), 1380–1387. 10.1038/nn.2662

Fedtsova, N., & Turner, E. E. (2001). Signals from the ventral midline and isthmus regulate the development of Brn3.0-expressing neurons in the midbrain. Mechanisms of development, 105(1-2), 129–144. 10.1016/s0925-4773(01)00399-9

Fujita, T., Epperly, M. W., Zou, H., Greenberger, J. S., & Wan, Y. (2008). Regulation of the anaphase-promoting complex-separase cascade by transforming growth factor-beta modulates mitotic progression in bone marrow stromal cells. Molecular biology of the cell, 19(12), 5446–5455. 10.1091/mbc.e08-03-0289

Furuta, Y., & Hogan, B. L. M.. (1998). BMP4 is essential for lens induction in the mouse embryo. Genes & Development, 12(23), 3764–3775. 10.1101/gad.12.23.3764

García-Campmany, L., & Martí, E. (2007). The TGFbeta intracellular effector Smad3 regulates neuronal differentiation and cell fate specification in the developing spinal cord. Development (Cambridge, England), 134(1), 65–75. 10.1242/dev.02702

Gray, G. E., & Sanes, J. R. (1992). Lineage of radial glia in the chicken optic tectum. Development (Cambridge, England), 114(1), 271–283. 10.1242/dev.114.1.271

Hamburger, V., & Hamilton, H. L. (1992). A series of normal stages in the development of the chick embryo. 1951. Developmental dynamics : an official publication of the American Association of Anatomists, 195(4), 231–272. 10.1002/aja.1001950404

Haramis, A. P., Begthel, H., van den Born, M., van Es, J., Jonkheer, S., Offerhaus, G. J., & Clevers, H. (2004). De novo crypt formation and juvenile polyposis on BMP inhibition in mouse intestine. Science (New York, N.Y.), 303(5664), 1684–1686. 10.1126/science.1093587

Hata, A., Lagna, G., Massagué, J., & Hemmati-Brivanlou, A. (1998). Smad6 inhibits BMP/Smad1 signaling by specifically competing with the Smad4 tumor suppressor. Genes & development, 12(2), 186–197. 10.1101/gad.12.2.186

Hébert, J. M., Mishina, Y., & Mcconnell, S. K.. (2002). BMP Signaling Is Required Locally to Pattern the Dorsal Telencephalic Midline. Neuron, 35(6), 1029–1041. 10.1016/s0896-6273(02)00900-5

Hegarty, S. V., Collins, L. M., Gavin, A. M., Roche, S. L., Wyatt, S. L., Sullivan, A. M., & O’Keeffe, G. W. (2014). Canonical BMP-Smad signalling promotes neurite growth in rat midbrain dopaminergic neurons. Neuromolecular medicine, 16(2), 473–489. 10.1007/s12017-014-8299-5

Hegarty, S. V., O’Keeffe, G. W., & Sullivan, A. M. (2013). BMP-Smad 1/5/8 signalling in the development of the nervous system. Progress in neurobiology, 109, 28–41. 10.1016/j.pneurobio.2013.07.002

Huber, C., Anand, A. A., Mauz, M., Künstle, P., Hupp, W., Hirt, B., & Wizenmann, A. (2013). In ovo expression of microRNA in ventral chick midbrain. Journal of visualized experiments : JoVE, (79), e50024. 10.3791/50024

Ille, F., Atanasoski, S., Falk, S., Ittner, L. M., Märki, D., Büchmann-Møller, S., Wurdak, H., Suter, U., Taketo, M. M., & Sommer, L. (2007). Wnt/BMP signal integration regulates the balance between proliferation and differentiation of neuroepithelial cells in the dorsal spinal cord. Developmental biology, 304(1), 394–408. 10.1016/j.ydbio.2006.12.045

Itasaki, N., Bel-Vialar, S., & Krumlauf, R. (1999). ’Shocking’ developments in chick embryology: electroporation and in ovo gene expression. Nature cell biology, 1(8), E203–E207. 10.1038/70231

Itoh, S., & Ten Dijke, P. (2007). Negative regulation of TGF-β receptor/Smad signal transduction. Current Opinion in Cell Biology, 19(2), 176–184. 10.1016/j.ceb.2007.02.015

Jardin, N., Giudicelli, F., Martín, D. T., Vitrac, A., Gois, S. D., Allison, R., Houart, C., Reid, E., Hazan, J., & Fassier, C. (2018). BMP- and neuropilin 1-mediated motor axon navigation relies on spastin alternative translation. Development (Cambridge, England), 145(17), dev162701. 10.1242/dev.162701

Jovanovic, V. M., Salti, A., Tilleman, H., Zega, K., Jukic, M. M., Zou, H., Friedel, R. H., Prakash, N., Blaess, S., Edenhofer, F., & Brodski, C. (2018). BMP/SMAD Pathway Promotes Neurogenesis of Midbrain Dopaminergic Neurons *In Vivo* and in Human Induced Pluripotent and Neural Stem Cells. The Journal of neuroscience : the official journal of the Society for Neuroscience, 38(7), 1662–1676. 10.1523/JNEUROSCI.1540-17.2018

Kitisin, K., Saha, T., Blake, T., Golestaneh, N., Deng, M., Kim, C., Tang, Y., Shetty, K., Mishra, B., & Mishra, L. (2007). Tgf-Beta signaling in development. Science’s STKE : signal transduction knowledge environment, 2007(399), cm1. 10.1126/stke.3992007cm1

Lee, K. J., Dietrich, P., & Jessell, T. M. (2000). Genetic ablation reveals that the roof plate is essential for dorsal interneuron specification. Nature, 403(6771), 734–740. 10.1038/35001507

Lee, K. J., Mendelsohn, M., & Jessell, T. M. (1998). Neuronal patterning by BMPs: A requirement for GDF7 in the generation of a discrete class of commissural interneurons in the mouse spinal cord. Genes & Development, 12(21), 3394. 10.1101/gad.12.21.3394

Li, N., Hornbruch, A., Klafke, R., Katzenberger, B., & Wizenmann, A. (2005). Specification of dorsoventral polarity in the embryonic chick mesencephalon and its presumptive role in midbrain morphogenesis. Developmental dynamics : an official publication of the American Association of Anatomists, 233(3), 907–920. 10.1002/dvdy.20434

Liem, K. F., Jr, Tremml, G., & Jessell, T. M. (1997). A role for the roof plate and its resident TGFbeta-related proteins in neuronal patterning in the dorsal spinal cord. Cell, 91(1), 127–138. 10.1016/s0092-8674(01)80015-5

Liem, K. F., Jr, Tremml, G., Roelink, H., & Jessell, T. M. (1995). Dorsal differentiation of neural plate cells induced by BMP-mediated signals from epidermal ectoderm. Cell, 82(6), 969–979. 10.1016/0092-8674(95)90276-7

Lipovsek, M., Ledderose, J., Butts, T., Lafont, T., Kiecker, C., Wizenmann, A., & Graham, A. (2017). The emergence of mesencephalic trigeminal neurons. Neural development, 12(1), 11. 10.1186/s13064-017-0088-z

Lo, L., Dormand, E. L., & Anderson, D. J. (2005). Late-emigrating neural crest cells in the roof plate are restricted to a sensory fate by GDF7. Proceedings of the National Academy of Sciences, 102(20), 7192–7197. 10.1073/pnas.0502581102

Martínez, V. G., Sacedón, R., Hidalgo, L., Valencia, J., Fernández-Sevilla, L. M., Hernández-López, C., Vicente, A., & Varas, A. (2015). The BMP Pathway Participates in Human Naive CD4+ T Cell Activation and Homeostasis. PLOS ONE, 10(6), e0131453. 10.1371/journal.pone.0131453

Millonig, J. H., Millen, K. J., & Hatten, M. E. (2000). The mouse Dreher gene Lmx1a controls formation of the roof plate in the vertebrate CNS. Nature, 403(6771), 764–769. 10.1038/35001573

Miyazono, K., Kamiya, Y., & Morikawa, M. (2010). Bone morphogenetic protein receptors and signal transduction. Journal of biochemistry, 147(1), 35–51. 10.1093/jb/mvp148

Momose, T., Tonegawa, A., Takeuchi, J., Ogawa, H., Umesono, K., & Yasuda, K. (1999). Efficient targeting of gene expression in chick embryos by microelectroporation. Development, growth & differentiation, 41(3), 335–344. 10.1046/j.1440-169x.1999.413437.x

Nguyen, V. H., Trout, J., Connors, S. A., Andermann, P., Weinberg, E., & Mullins, M. C. (2000). Dorsal and intermediate neuronal cell types of the spinal cord are established by a BMP signaling pathway. Development (Cambridge, England), 127(6), 1209–1220. 10.1242/dev.127.6.1209

Nohe, A., Keating, E., Knaus, P., & Petersen, N. O. (2004). Signal transduction of bone morphogenetic protein receptors. Cellular signalling, 16(3), 291–299. 10.1016/j.cellsig.2003.08.011

Hegarty, S. V., & Sullivan, A. M. (2017). Targeting bone morphogenetic protein signalling in midbrain dopaminergic neurons as a therapeutic approach in Parkinson’s disease. Neuronal Signaling, 1(2), NS20170027. 10.1042/NS20170027

Patten, I., & Placzek, M. (2002). Opponent activities of Shh and BMP signaling during floor plate induction in vivo. Current biology : CB, 12(1), 47–52. 10.1016/s0960-9822(01)00631-5

Piccolo, S., Sasai, Y., Lu, B., & De Robertis, E. M. (1996). Dorsoventral patterning in Xenopus: inhibition of ventral signals by direct binding of chordin to BMP-4. Cell, 86(4), 589–598. 10.1016/s0092-8674(00)80132-4

Quinlan, R., Graf, M., Mason, I., Lumsden, A., & Kiecker, C. (2009). Complex and dynamic patterns of Wnt pathway gene expression in the developing chick forebrain. Neural development, 4, 35. 10.1186/1749-8104-4-35

Roussa, E., von Bohlen und Halbach, O., & Krieglstein, K. (2009). TGF-beta in dopamine neuron development, maintenance and neuroprotection. Advances in experimental medicine and biology, 651, 81–90. 10.1007/978-1-4419-0322-8_8

Seoane, J., Pouponnot, C., Staller, P., Schader, M., Eilers, M., & Massagué, J. (2001). TGFbeta influences Myc, Miz-1 and Smad to control the CDK inhibitor p15INK4b. Nature cell biology, 3(4), 400–408. 10.1038/35070086

Smidt, M. P., & Burbach, J. P. (2007). How to make a mesodiencephalic dopaminergic neuron. Nature Reviews Neuroscience, 8(1), 21–32. 10.1038/nrn2039

Solloway, M. J., & Robertson, E. J. (1999). Early embryonic lethality in Bmp5;Bmp7 double mutant mice suggests functional redundancy within the 60A subgroup. Development (Cambridge, England), 126(8), 1753–1768. 10.1242/dev.126.8.1753

Timmer, J. R., Wang, C., & Niswander, L. (2002). BMP signaling patterns the dorsal and intermediate neural tube via regulation of homeobox and helix-loop-helix transcription factors. Development (Cambridge, England), 129(10), 2459–2472. 10.1242/dev.129.10.2459

Tsang, H. T., Edwards, T. L., Wang, X., Connell, J. W., Davies, R. J., Durrington, H. J., O’Kane, C. J., Luzio, J. P., & Reid, E. (2009). The hereditary spastic paraplegia proteins NIPA1, spastin and spartin are inhibitors of mammalian BMP signalling. Human molecular genetics, 18(20), 3805–3821. 10.1093/hmg/ddp324

Vonica, A., & Brivanlou, A. H. (2006). An obligatory caravanserai stop on the silk road to neural induction: inhibition of BMP/GDF signaling. Seminars in cell & developmental biology, 17(1), 117–132. 10.1016/j.semcdb.2005.11.013

Wang, X., Shaw, W. R., Tsang, H. T., Reid, E., & O’Kane, C. J. (2007). Drosophila spichthyin inhibits BMP signaling and regulates synaptic growth and axonal microtubules. Nature neuroscience, 10(2), 177–185. 10.1038/nn1841

Watanabe, Y., & Nakamura, H. (2000). Control of chick tectum territory along dorsoventral axis by Sonic hedgehog. Development (Cambridge, England), 127(5), 1131–1140. 10.1242/dev.127.5.1131

Watanabe, Y., & Yaginuma, H. (2015). Tangential cell migration during layer formation of chick optic tectum. Development, growth & differentiation, 57(8), 539–543. 10.1111/dgd.12238

Wilson, L., & Maden, M. (2005). The mechanisms of dorsoventral patterning in the vertebrate neural tube. Developmental biology, 282(1), 1–13. 10.1016/j.ydbio.2005.02.027

Wine-Lee, L., Ahn, K. J., Richardson, R. D., Mishina, Y., Lyons, K. M., & Crenshaw, E. B., 3rd (2004). Signaling through BMP type 1 receptors is required for development of interneuron cell types in the dorsal spinal cord. Development (Cambridge, England), 131(21), 5393–5403. 10.1242/dev.01379

Wozney, J. M., Rosen, V., Celeste, A. J., Mitsock, L. M., Whitters, M. J., Kriz, R. W., Hewick, R. M., & Wang, E. A. (1988). Novel regulators of bone formation: molecular clones and activities. Science (New York, N.Y.), 242(4885), 1528–1534. 10.1126/science.3201241

Xiao, Y. T., Xiang, L. X., & Shao, J. Z. (2007). Bone morphogenetic protein. Biochemical and biophysical research communications, 362(3), 550–553. 10.1016/j.bbrc.2007.08.045

Yan, H., Zhu, S., Song, C., Liu, N., & Kang, J. (2012). Bone morphogenetic protein (BMP) signaling regulates mitotic checkpoint protein levels in human breast cancer cells. Cellular signalling, 24(4), 961–968. 10.1016/j.cellsig.2011.12.019

Zhong, J., & Zou, H. (2014). BMP signaling in axon regeneration. Current opinion in neurobiology, 27, 127–134. 10.1016/j.conb.2014.03.009Zhong, J., Zou, H., 2014. BMP signaling in axon regeneration. Curr Opin Neurobiol 27, 127–134

